## Supplementary Information for "Connecting MHC-I-binding motifs with HLA alleles via deep learning"

Supplementary Table 1. Summary of the datasets used in training, validating, and evaluating the MHCfovea’s predictor

|  | D-E ratio | Binding Assay | | Ligand Elution | Protein Decoy | Random Decoy |
| --- | --- | --- | --- | --- | --- | --- |
|  |  | Positive | Negative |  |  |  |
| Training | 30 | 38,793 | 110,209 | 226,800 | 1,068,270 | 5,801,583 |
|  | 60 |  |  |  | 2,136,540 | 11,603,163 |
|  | 90 |  |  |  | 3,204,791 | 17,404,743 |
| Validation | 30 | 2,088 | 5,754 | 11,937 | 174,361 | 179,055 |
| Benchmark | 30 | 0 | 0 | 127,371 | 1,980,947 | 2,036,733 |

Supplementary Table 2. Data number of each dataset by alleles (Additional File)

The number of peptides by alleles from different sources including binding assay data, ligand elution data, and decoy data in the training, validation, and benchmark datasets.

Supplementary Table 3. Performance metrics by alleles of each predictor (Additional File)

The performance metrics including AUC, AP, AUC0.1, and PPV by alleles of different predictors. There are two kinds of results corresponding to comparison with MixMHCpred2.1 or without MixMHCpred2.1.

Supplementary Table 4. The list of the rare alleles in the benchmark dataset

| A*02:05 | A*34:02 | B*13:02 | B*40:06 | C*03:02 |
| --- | --- | --- | --- | --- |
| A*11:02 | A*36:01 | B*15:10 | B*52:01 | C*04:03 |
| A*24:07 | A*74:01 | B*35:07 | B*55:01 | C*07:04 |
| A*33:03 | B*07:04 | B*37:01 | B*55:02 | C*08:01 |
| A*34:01 | B*13:01 | B*38:02 | B*58:02 | C*14:03 |

Supplementary Table 5. Positional annotation of the MHC-I sequence (Additional File)

The annotation of the important positions of each HLA gene, the polymorphic positions, and the pseudo-sequence (34 a.a.) of NetMHCpan4.1.

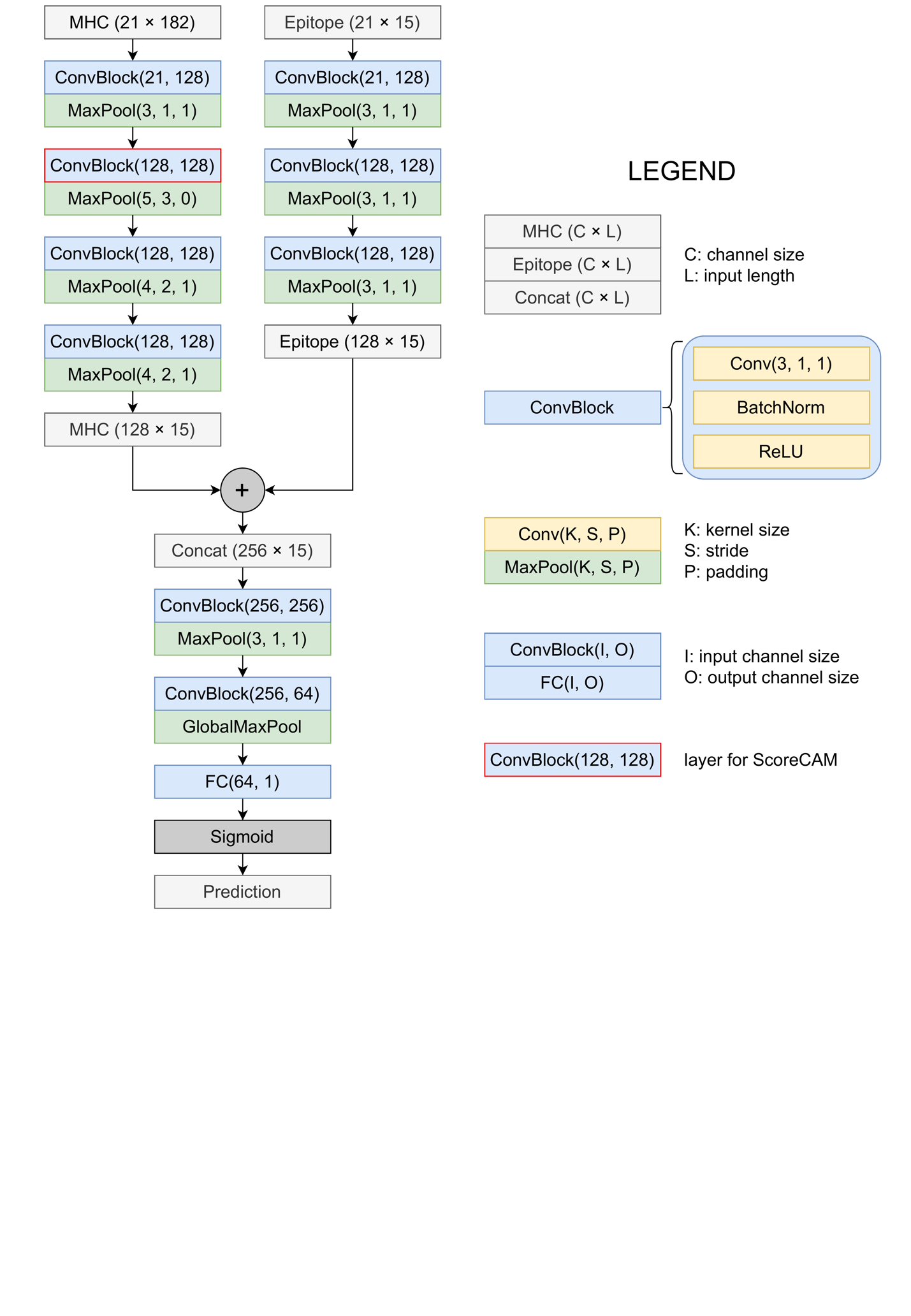
**Supplementary Figure 1. The CNN model architecture of MHCfovea’s predictor.** The predictor adopted by MHCfovea is an ensemble model of multiple CNN model. Each CNN model takes both MHC-I sequence and epitope sequence as input. The MHC part and epitope part pass through four and three convolution blocks separately. Then, they are concatenated on the dimension of convolutional channels, and pass through another two convolution blocks followed by a max global pooling layer, a fully connective layer, and a sigmoid function to get the final prediction score. The convolution block with a red box is the layer for the ScoreCAM process.

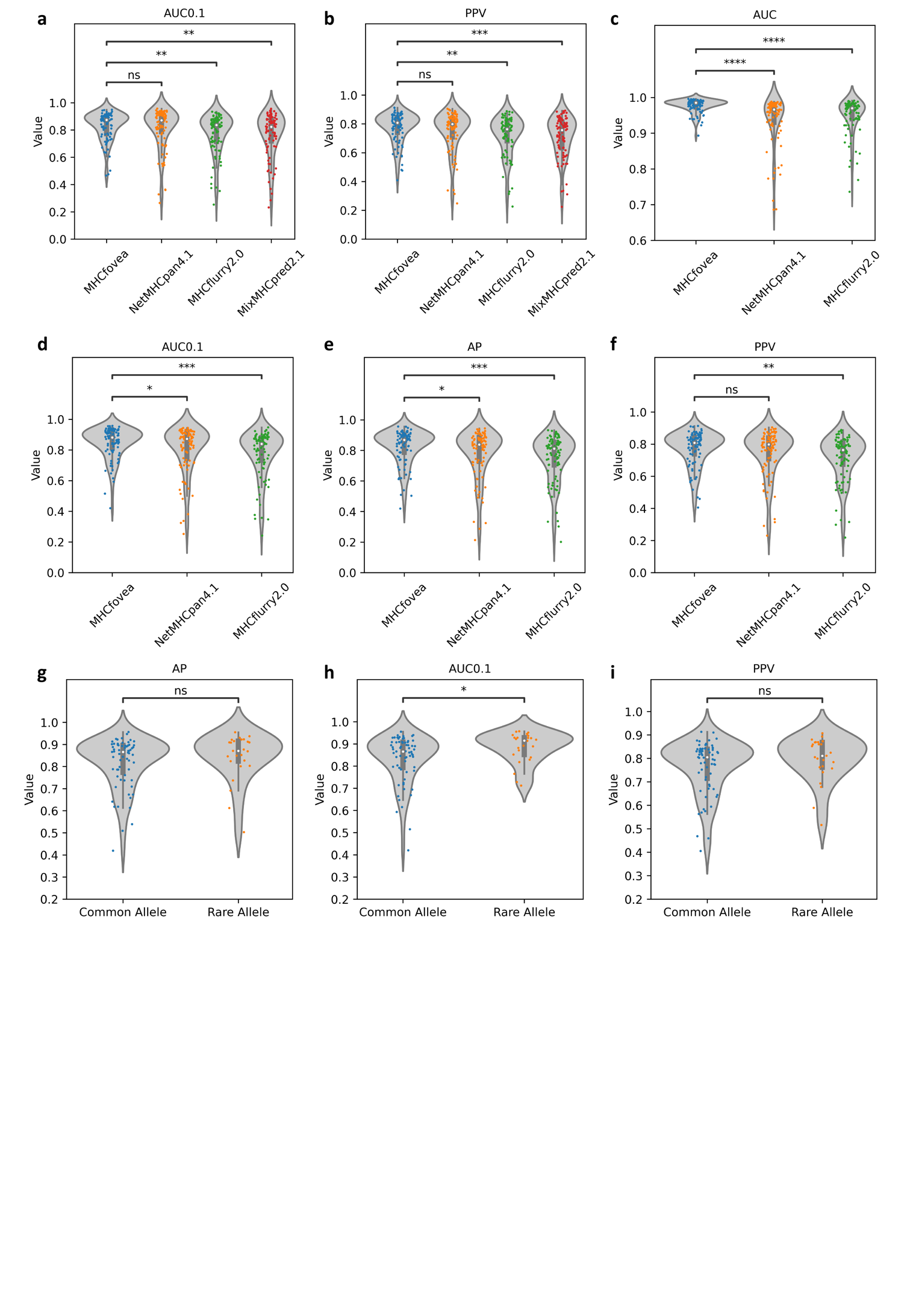
**Supplementary Figure 2. Performance of MHCfovea’s predictor.** Four metrics are used to evaluate the model performance, including AUC, AP, AUC0.1, and PPV. **a-b**, Violin plots depict the distribution of AUC0.1 and PPV on the benchmark dataset with peptides available for MixMHCpred2.1, where peptides beyond 20 amino acids and more than 14 amino acids are dropped (allele number = 91). AUC and AP are presented on Fig. 2d, 2e. **c-f**, Comparison of four metrics between MHCfovea, NetMHCpan4.1, and MHCflurry2.0 on the full benchmark dataset (allele number = 92). **g-i**, The distribution of AP, AUC0.1, and PPV on common alleles (n = 67) and 25 rare alleles. The AUC distribution is shown in Fig. 2f. Boxplots depict the median value with a white dot, the 75^th^ and 25^th^ percentile upper and lower hinges, respectively, and whiskers with 1.5x interquartile ranges. P-values (two-tailed independent t-test) are shown as “ns” no significance, * P ≤ 0.05, ** P ≤ 0.01, *** P ≤ 0.001, and **** P ≤ 0.0001.

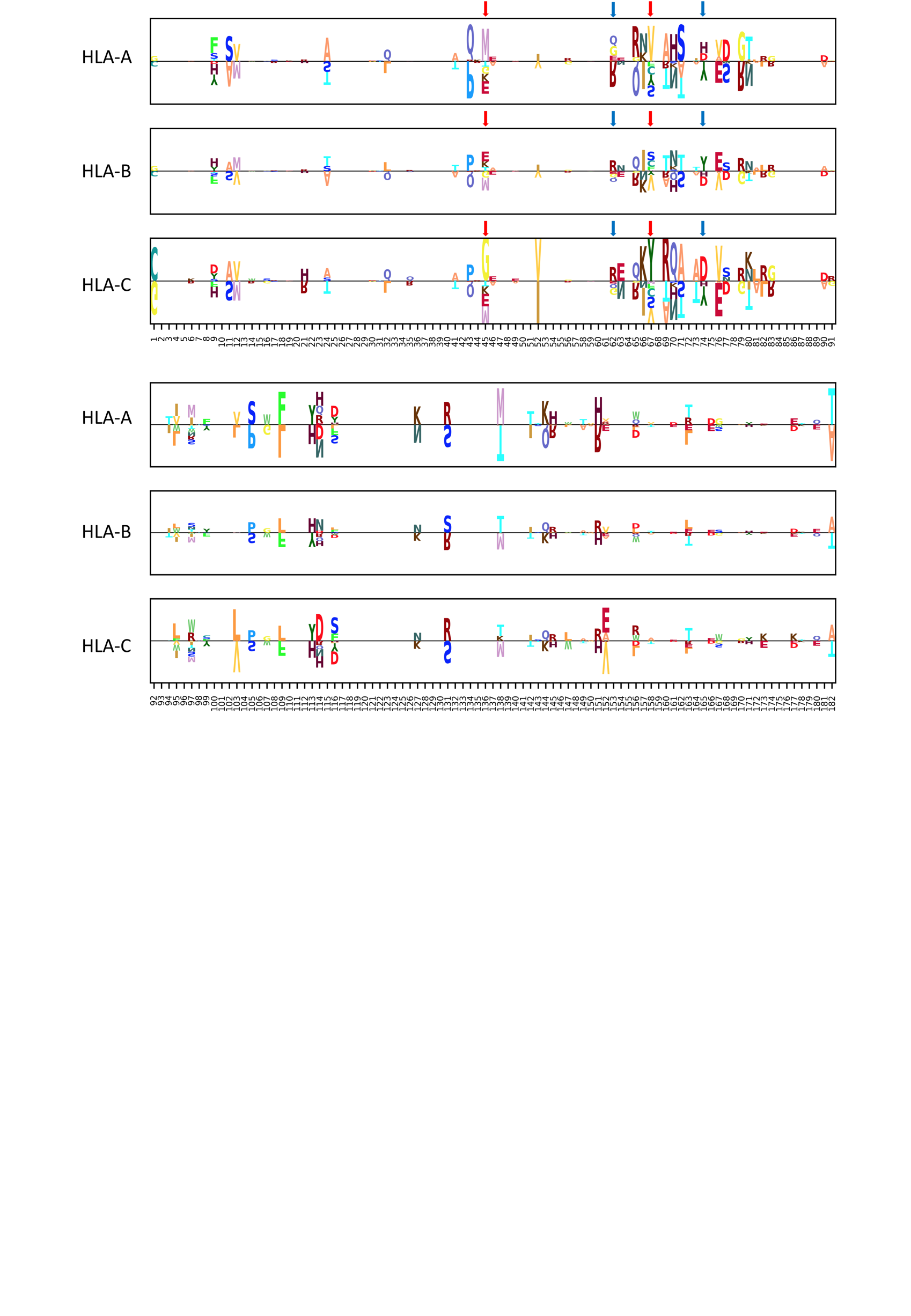
**Supplementary Figure 3. The sequence logo of MHC-I sequence derived from different HLA genes.** Only alleles from the training dataset are used to construct the position probability matrix (PPM). Each sequence logo is the difference between PPM of a specific HLA gene and PPM of all alleles. Each HLA gene has its own patterns on particular regions. Some positions are highly polymorphic for an HLA gene but conservative for others. For example, the positions annotated by red arrows are polymorphic in HLA-B but conservative for HLA-A and -C. In the same way, the positions annotated by blue arrows are polymorphic in HLA-A but conservative for HLA-B and -C.

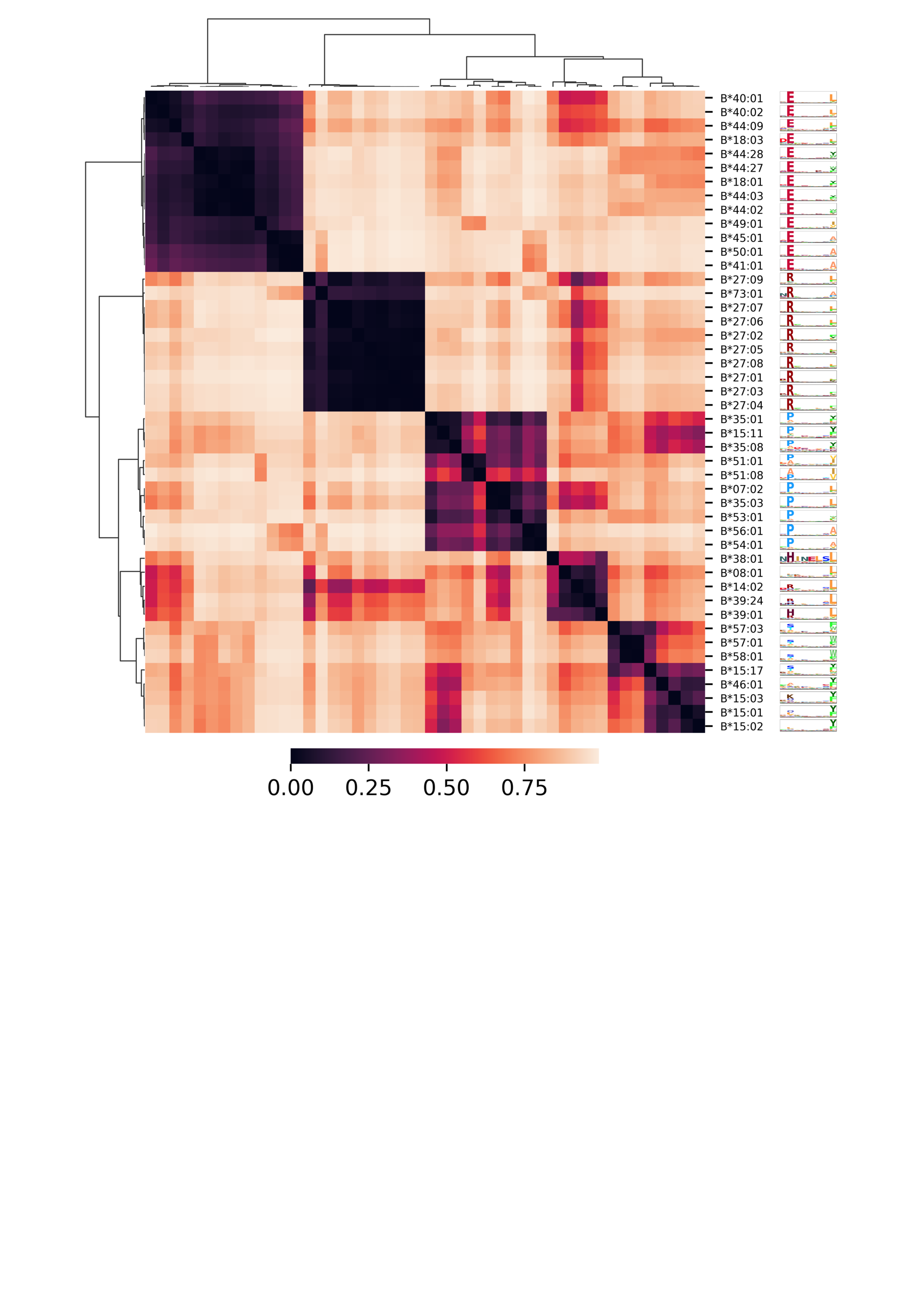
**Supplementary Figure 4. The heatmap clustering of MHC-I-binding motifs of HLA-B alleles in the training dataset.** A hierarchical clustering with cosine distance metric and UPGMA (unweighted pair group method with arithmetic mean) algorithm is used. In this figure, the clustering result was either dominated by the N-terminal or the C-terminal sub-motifs of the alleles. Alleles with similar N-terminal sub-motifs may have dissimilar C-terminal sub-motifs. For example, both HLA-B*07:02 and HLA-B*56:01 have a P-dominant N-terminal sub-motif, but the former has an L-dominant C-terminal sub-motif and the latter has an A-dominant one.

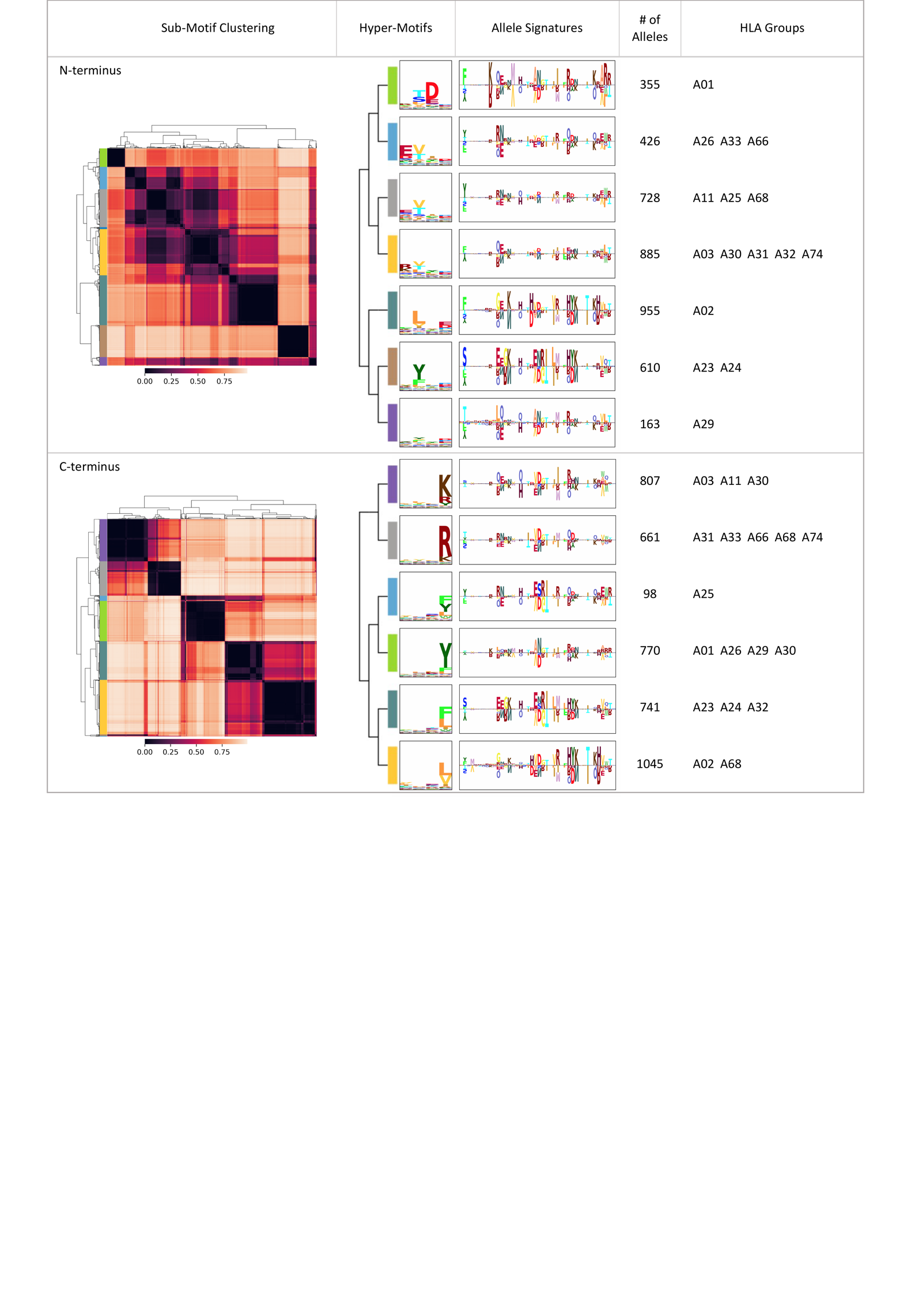
**Supplementary Figure 5. A summarization table of HLA-A.** The MHC-I-binding motifs are divided into N-terminal and C-terminal sub-motifs; sub-motifs are clustered by agglomerative hierarchical clustering. Hyper-motifs and the corresponding allele signatures are calculated for each sub-motif cluster. In each cluster, the number of alleles, and the HLA groups with the number of alleles ≥25, are recorded in the last two columns.

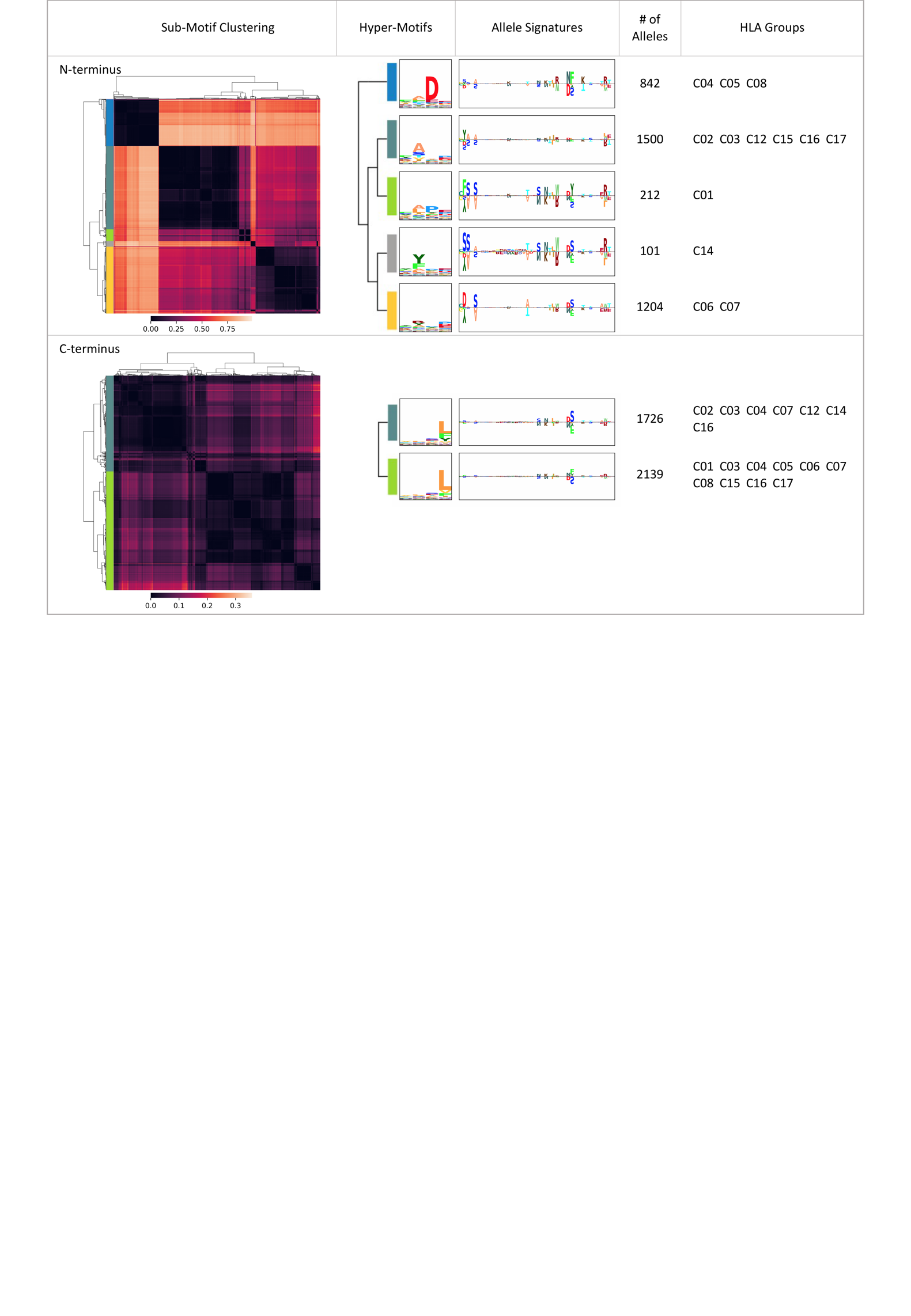
**Supplementary Figure 6. A summarization table of HLA-C.** The MHC-I-binding motifs are divided into N-terminal and C-terminal sub-motifs; sub-motifs are clustered by agglomerative hierarchical clustering. Hyper-motifs and the corresponding allele signatures are calculated for each sub-motif cluster. In each cluster, the number of alleles, and the HLA groups with the number of alleles ≥25, are recorded in the last two columns.

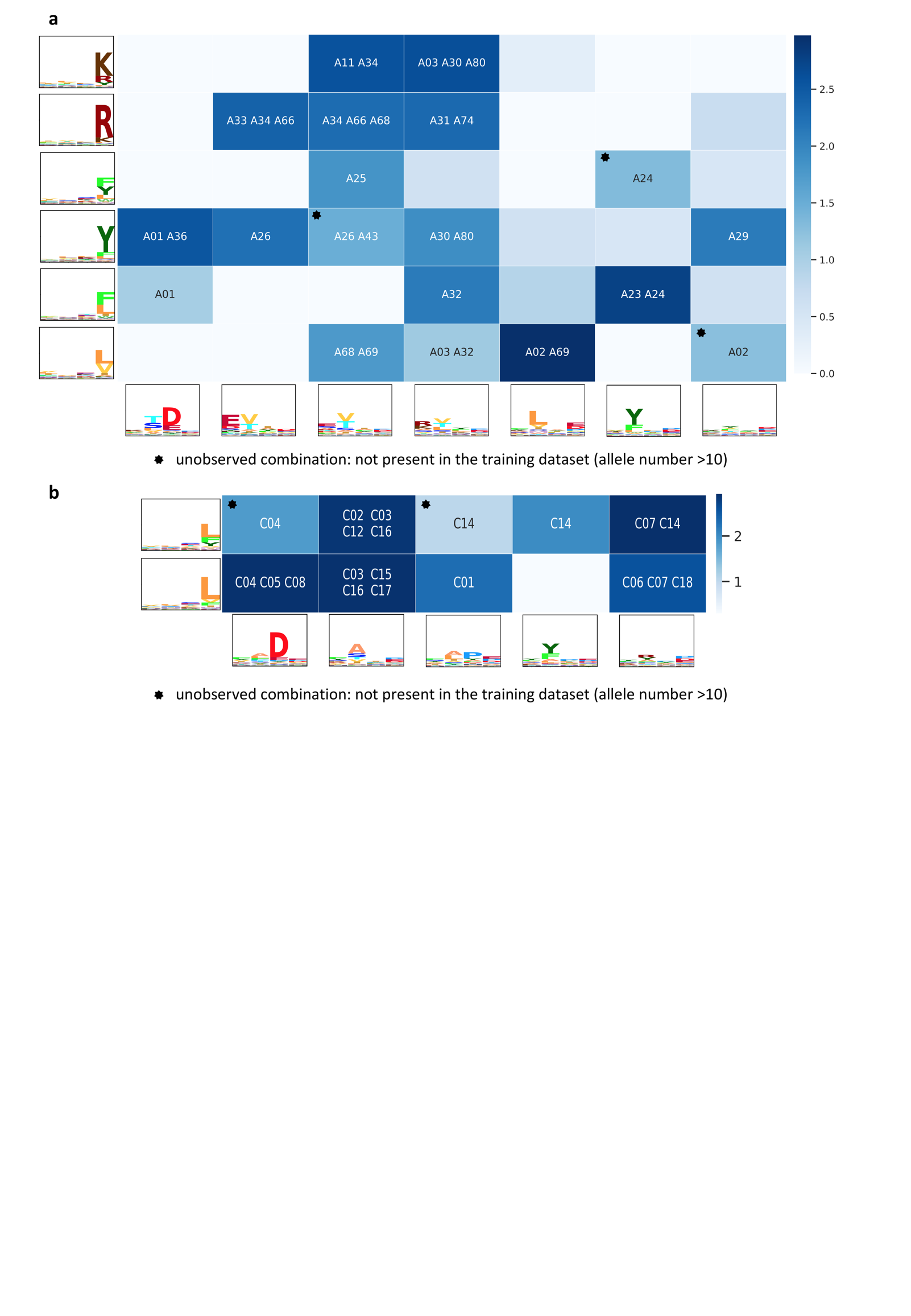
**Supplementary Figure 7. The combination map of N-terminal and C-terminal hyper-motifs. a-b**, The heatmap on the combinations of N-terminal (x-axis) and C-terminal (y-axis) hyper-motifs for HLA-A (**a**) and -C (**b**). The binding motif of an allele is a combination of an N-terminal and a C-terminal hyper-motif. After allocating all the alleles into the combination map, the cell color is determined by log_10_(number of alleles in the cell). In each cell with an allele number >10, the maximal HLA group, and HLA groups with an allele number ≥25, or with a proportion (the allele number in the cell to the overall number of an allele group) >0.1, are listed.
